# Development and evaluation of screwworm, *Cochliomyia hominivorax*, transgenic sexing strains with embryonic gene promoters for a genetic control program

**DOI:** 10.1101/2025.05.01.651745

**Authors:** Alex P. Arp, Aidamalia Vargas Lowman, Carolina Concha, Ying Yan, Andrea Martínez, Gladys Quintero, Mario Vazquez, Agustin Sagel, Maxwell J. Scott

**Affiliations:** USDA-ARS, Knipling-Bushland US Livestock Insects Research Laboratory, Kerrville, TX, 78216; USDA-ARS, Screwworm Research Laboratory, Pacora, Panama; Panama-United States Commision for the Eradication and Prevention of Screwworm (COPEG), Pacora, Panama; Justus-Liebig-University Giessen, Institute for Insect Biotechnology, Department of Insect Biotechnology in Plant Protection, Winchesterstraße 2, 35394 Gießen, Germany; Escuela de Biotecnología. Facultad de Ciencias de la Salud, Universidad Latina de Panamá, Panama City, Panama; Department of Entomology and Plant Pathology, North Carolina State University, Raleigh, NC, 27695, USA; Department of Genetics and Molecular Biology, University of Panama, Panama City, Panama. Member of SNI-SENACYT Panama

**Keywords:** sexing strain, sterile insect technique, genetic biocontrol, New World screwworm

## Abstract

The New World screwworm, *Cochliomyia hominivorax*, was eradicated from North and Central America through the first application of the sterile insect technique. The sterile screwworm adult fly releases were mixed sex, but experience with other flies suggests that releasing only males could be up to five times more effective. Here we describe screwworm transgenic sexing strains (TSSs) with expected embryo lethality developed using the Tetracycline-off (Tet-off) system with the *Lucilia cuprina nullo* (DR6) or *C. macellaria* CG14427 (DR7) gene promoters and a tTA activated effector to promote female lethality. The TSSs expressed tTA highest in 2-3 h embryos and low in larvae and adults. However, most strains showed high expression in pupae. Evaluation on two doxycycline (Dox) regimens found that inclusion of Dox in the last larval feeding rescued females of the subsequent generation, likely by maternal Dox transfer to embryos. All TSSs produced only males on a reduced Dox feeding regimen but the female lethal period for the DR6 TSS was too late in development to save diet costs. Production parameters were met by all strains in colony, but strains had lower male fly survival than current production strains after removing Dox. In non-competitive mating success trials DR6 strains performed poorly, but DR7 performed equally to production strain males. However, males from all TSSs faired poorly in mating competition tests against production males. Our study highlights the importance of tightly-regulated gene promoters and suitable antibotics feeding schemes for the development and evaluation of TSSs based on Tet-off system.

## Introduction

The New World screwworm, *Cochliomyia hominivorax*, is a blowfly with an obligate parasitic larval stage that uses warm blooded animals as hosts (Pereira de Barros and Bricarello 2020). Developing larvae are covered in spines and their constant screwing motion causes significant damage to host tissues, often leading to death if untreated. Screwworms are most prevalent in livestock but are generalists and will infest all warm-blooded animals including humans (Pereira de Barros and Bricarello 2020, Batista-da-Silva et al. 2011,). Control for screwworm is primarily reliant on limiting birthing or farming practices that result in wounds to seasons of low screwworm presence and extensive surveillance (Novy 1991). In addition to cultural controls, the sterile insect technique (SIT) was developed for screwworm and used to eradicate the species from all North and Central America between the 1950’s and 2000’s (Knipling 1985; Vargas-Terán et al. 2020). From 2006 a barrier zone was established on the Panama-Colombia border operated by the Panama-United States Commission for the Eradication and Prevention of Screwworm (COPEG). To prevent the fly from returning from South America, about 15 million sterile males and females were released weekly in the barrier zone. However, in 2023 the barrier failed due to unknown reasons and screwworm rapidly spread throughout Central America and entered Mexico in December 2024 (USDA APHIS, https://www.aphis.usda.gov/livestock-poultry-disease/cattle/ticks/screwworm/outbreak-central-america). The breakdown of the screwworm SIT program highlights the need to continuously develop novel control strategies and technology for pest control.

The efficacy of SIT programs relies on their ability to produce highly fit males able to outcompete or overwhelm wild males to mate fertile females. It was predicted by Herman Muller in correspondence with E.F. Knipling that genetic suppression would be improved if sterile females are excluded from releases since they distract sterile males and do not contribute to genetic suppression (Muller 1950, Letter to Knipling). Indeed, male-only field releases of sterile Medfly, *Ceratitis capitata*, was several-fold more effective than bisexual releases (Hendrichs et al. 2003; Rendón et al. 2004). Since sterile females are not considered effective for reducing wild populations, their removal early in development can infer large cost savings for production through the reduction of diet material and labor (Chen et al. 2014; Concha et al. 2020). The screwworm eradication program has operated since inception using mixed-sex sterile releases due to the lack of dimorphic features in pupae or larvae. Additionally, classical genetic sexing strains, such as those used by the Medfly SIT programs have not been developed for screwworm (Caceres 2002).

To produce male-only screwworm strains, genetic engineering technologies were used. The first transgenic screwworm strains were created with *piggyBac* (*pBac*) transposase and expressed a fluorescent marker gene proving that transgenic strain development was a possibility for this species (Allen et al. 2004). Transgenic sexing strains (TSSs) of screwworm were then developed utilizing a tetracycline-controlled transactivator (tTA) overexpression gene construct with the tTA gene modified to only express in females by addition of the *C. hominivorax transformer* (*Chtra*) sex-specific intron (Gong et al. 2005; Concha et al. 2016). The female *Chtra* intron contains an exon that is retained in males and contains in-frame stop codons (Li et al. 2013). Since the intron is spliced out by females, only females express tTA protein resulting in lethality (Concha et al. 2016). Overexpression of tTA is thought to cause lethality due a general interference in gene expression (i.e. “transcriptional squelching”) and/or interference with protein ubiquitination (Gong et al. 2005, Knudsen et al. 2020). Although these TSSs were female-lethal, females did not die until the pupal stage and did not confer any cost savings in comparison with production screwworm strains. Embryo-female lethal TSSs of screwworm were subsequently produced using a double-component driver and effector system (Ogaugwu et al. 2013; Concha et al. 2020). The driver construct utilized the *L. sericata bottleneck* (*Lsbnk*) promotor to drive production of tTA in early embryos when the *Lsbnk* gene is most active (Edman et al. 2015). The tTA protein then activated the *Lshid^Ala2^* proapoptotic gene which was driven by *tetO21-hsp70* enhancer-promoter. As the *Lshid^Ala2^* gene contained the *Chtra* sex-specific intron only female embryos died due to widespread induction of apoptosis. Although female mortality of strains carrying these constructs occurred in the embryo and first instar, tTA expression also occurred in developing female ovaries resulting in infertility in most, but not all, strains (Concha et al. 2020). Female fertility could be restored through providing tetracycline (Tc) in the water for 24 hours after emergence. Dosing females with Tc complicated the rearing process and posed a risk of maternal transfer of Tc to the offspring and rescuing female offspring (Schetelig et al. 2016, Yan et al. 2017, Yan et al. 2023). One strain, TD1#12, which has the driver and effector combined into a single all- in-one construct, did not require adult provisioning of Tc to maintain female fertility.

With the aim of obtaining early embryo-specific expression of tTA, drivers DR6 and DR7 were developed with promoters from *L. cuprina nullo* and *Cochliomyia macellaria* CG14427 genes, respectively (Yan et al. 2020). The *L. cuprina nullo* and *CG14427* genes showed higher expression in early embryos and lower expression in adult females than the *L. cuprina bottleneck* gene (Yan et al. 2020). DR6 and DR7 appeared to be an improvement over the DR2 driver as two component *L. cuprina* strains were found be embryonic lethal and there was no detectable tTA expression in ovaries making them more suitable for mass rearing. The aim of this study was to develop and evaluate DR6 and DR7 drivers in screwworm. Two component TSSs were evaluated for driver expression, survival, biological performance in accordance with mass-rearing parameters, mating capability, and longevity. As maternal transfer of Dox inhibited embryo lethality, we also evaluated the strains on a reduced Dox feeding regimen.

## Methods

### Insect rearing

Screwworm lines were reared in COPEG mass rearing facility in Pacora, Panama. The plant exceeds ACL-2 standards and development and testing of transgenic screwworm lines was approved by the Panamanian Biosecurity Commission.

Beginning with egg incubation and throughout larval development, larvae were held in shallow plastic food containers (12.5 cm x 20 cm x 4 cm) placed above approximately 2cm of sawdust lining the bottom of a larger tub (23 cm x 28 cm x 11 cm). From incubation to 120 hr, larvae were maintained at 37°C and 70% RH. To induce crawl off, larval trays were moved to a separate room maintained at 33°C and 50-60% RH, at 120 hr to 168 hr. Forty-eight hours after crawl-off, larvae were sifted from the sawdust and measured. Approximately 25 ml of pupae were randomly selected to produce the adult colony. All other pupae were used for additional assays or were discarded. Pupae and adults are held in the same room at 25.5°C and 55% RH under 12 hr:12 hr light:dark cycle.

Oviposition was induced using ∼10 g of raw ground beef scented with liquid from old larval diet on top of a plastic sauce cup filled with 45°C water.

Screwworm larvae were fed an artificial diet containing: 0.9% spray-dried bovine blood plasma (AP920, APC, Inc, Ankeny, IA, USA), 3.6% spray-dried bovine red blood cells (AP301, APC, Inc, Ankeny, IA, USA), 5.0% poultry egg powder (OVA-40, IsoNova Technologies, Edem Valley, MN, USA), 4.5% soy powder (Textured Soy Proteins, Archer-Daniels-Midland Company, Chicago, IL, USA), 0.0025% potassium permanganate (P279, Fisher Scientific, Fair Lawn, NJ, USA), 5% cellulose fiber (CF100, J Rettenmaier, Schoolcraft, MI, USA) and, if specified, 25 μg/ml doxycycline HCl (Laboratorio Hispanoamericano, S.A. (LHISA), La Libertad, El Salvador). Each test generation was seeded with 75 mg of eggs. Fresh diet was added daily in increasing increments (+Dox: 30 ml, 70 ml, 300 ml, 800 ml, -Dox: 30 ml, 45 ml, 150 ml, 400 ml) until crawl-off was induced on day 5.

Adults were fed a gelled diet containing 40% brown sugar (Santa Rosa, Azucarera Nacional, El Roble de Aguadulce, Cocle, Republic of Panama), 5% egg powder (OVA-40, IsoNova Technologies, Edem Valley, MN, USA), and 1% carrageenan (CEAMGEL 1725, CEAMSA, Pontevedra, Spain), and given fresh water through a cotton wick.

Doxycycline was added to the larval diet and adult water following two dosage regimens (Figure 1). In both regimens doxycycline was mixed in the diet at 25 μg/ml. In the full dosage regimen Dox was included in all Dox+ larval feedings and the adult water, while in the reduced Dox regimen Dox was only included in the first 3 larval feedings and adult water for Dox+ cages. In all Dox-treatments the parental generation were not given water with Dox prior to producing a Dox-generation.

**Figure 1.**
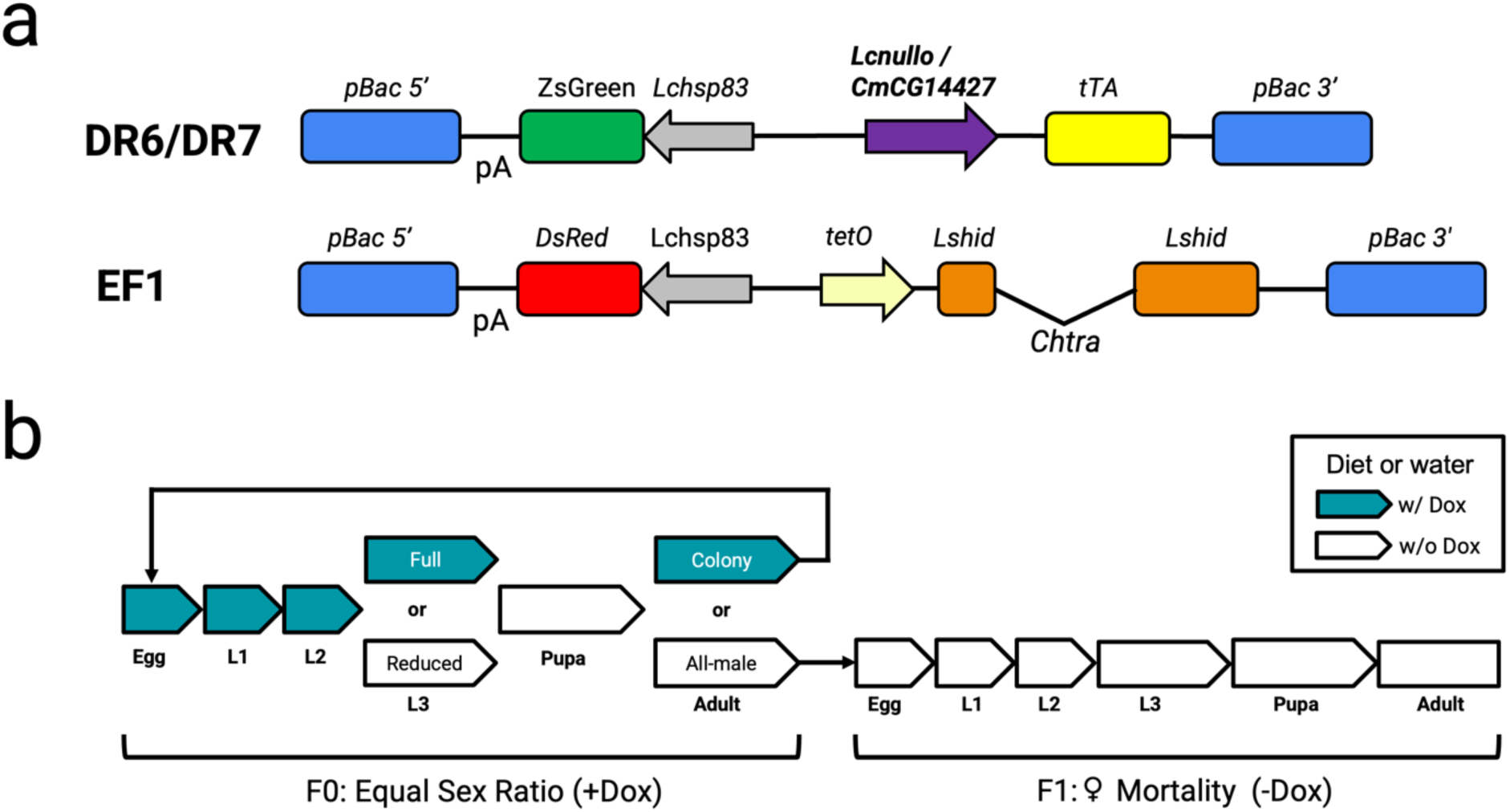
Two component transgenic sexing systems. a) Diagram of DR6, DR7 and EF1 gene constructs. DR6 contains the *Lcnullo* promoter while DR7 has the *CmCG14427* promoter to drive transcription of tTAo. Both DR6 and DR7 have a ZsGreen fluorescent marker driven by the *Lchsp83* promoter. The effector construct EF1 contains the tTA-inducible tetO-hsp70 enhancer-promoter to drive Lshid^Ala2^ production in females. b) Diagram of Dox feeding regimes used to maintain DR6/DR7 x EF1 strains and produce male-only generations. Strains were evaluated on a full dosage regimen with all larval diet additions containing Dox, or a reduced Dox regimen where the final larval diet addition did not contain Dox. Adults used for colony maintenance were provided Dox in their water, while adults used for producing an all-male generation were not provided Dox. L1, L2, L3 indicate larval instars.

### Gene constructs

The construction of the DR6 and DR7 plasmids was described previously (Yan et al. 2020). The plasmid sequences are available from NCBI under the accession numbers MK509802 for DR6 and MK509803 for DR7.

### Strain creation

Strains were developed using the screwworm mass-rearing strain Jamaica-06 Crio 13 (J-06). Six-day old females were induced to oviposit on the lid of a 2 oz sauce cup filled with 40°C water and topped with raw ground beef scented with liquid from spent larval diet media. Cages were checked every 10-15 minutes for fresh eggs. Eggs were removed and washed in 2% KOH solution for 2 min to separate egg masses then rinsed with DI water. Eggs were then aligned on double sided tape on a welled microscope slide. Slides were desiccated for 6 min in a chamber with silica desiccant beads. After desiccation eggs were covered with Mineral Oil 27 (M8410, Sigma-Aldrich, St. Louis, MO, USA) prior to injection.

Injections were completed using a Sutter XenoWorks Digital Injector and Micromanipulator (Sutter Instrument, Novato, CA) with a compound microscope. Needles used for injection were made from filamented Quartz glass capillary tubes (QF100-70-10; Sutter Instruments, Novato, CA) pulled on a Sutter P-2000 (Program, Heat: 650, Fill: 04, Velocity: 50, Delay: 150, Pull: 155). Embryos were injected with a mix of transgene vector plasmid, pBac expressing plasmid, and pBac mRNA as previously described (Concha et al. 2016). After injection oil was removed and slides were placed in an oxygen rich chamber in an incubator at 37°C overnight. The following day transiently expressing G0 larvae were collected under a fluorescent stereoscope and reared to adult. G0 males were individually crossed to 5 virgin wild type J-06 females, and G0 females were placed with two J-06 males. G1 offspring were then screened for fluorescence. Any expressing G1 offspring were then individually crossed to wild type J-06 to verify single insertion and heritable transgene integration. One G1 from each G0 was used to produce a stable strain by inbreeding and filtering for homozygosity from G2 onward.

### Expression analysis

Whole insect samples were frozen at -80°C prior to RNA extraction. Total RNA was isolated from samples using a Zymo Direct-zol MiniPrep kit (R2051, Zymo Research Group, Irvine, CA, USA) according to manufacturer recommendations. After extraction, RNA concentration was assessed using a NanoDrop 1000. cDNA libraries were created using a Aquaris qMax kit (PR 2100-C, Accuris Instruments, Edison, NJ, USA) using 50 ng of total RNA to produce 10 μl of cDNA solution. cDNA products were diluted to 1/20 concentration for use in PCR analysis.

Expression of tTAo was assessed using GAPDH as the reference gene using the following primers: GAPDH-F: GTCAGTGACACCCACTCCTC, GAPDH-R: TTGATCAAGTCGATGACACG, tTAo-F: TCTTGCGTAATAATGCCAAATCCTTCCG, tTAo-R: CCAACACACAGCCCAATGTAAAATGACC (Cardoso et al. 2014, Li et al. 2014). PCR was done on a Mic qPCR Cycler (Bio Molecular System, Australia) with 10 μl reactions using AB Power Up SYBR green master mix (A25780, Life Technologies, Burlington, ONT, Canada). PCR cycles as follows: Activation at 50°C 2 min followed by 95°C for 2 min; Cycle 40 times of 95°C for 3 sec and 60°C for 30 sec.

Expression ratios were evaluated using the ΔΔCt method with the 2^nd^ instar larvae sample expression used as the calibrator. Fifteen grams of eggs were collected at precellular (0-1 hr), cellular (2-3 hr) and late stage embryos (5-6 hr). As well, as samples of second instar larvae at 48 hr, pupae of 48 hr and young adult female of 24 hr. Expression was evaluated from three separate samples for each stage and each reaction was run in triplicate.

### Strain performance evaluation

Egg hatch rate was evaluated by placing 200 individual embryos on a wet circular 95 mm diameter black filter paper inside a petri dish and incubating for 24 hr at 37°C. Empty eggs were then counted as hatched. Two trays per cage were counted every generation.

Larval stage mortality was evaluated by counting larvae surviving from a 5mg mass of eggs. Three eggs masses were prepared for each cage and counted only once, either after 24 hr, 48 hr, or 72 hr. Eggs were placed on 5 mg of larval diet and an additional 10 ml were added at 24 hr and 50 ml at 48 hr. Larvae were held in the same conditions and fed the same diet as the corresponding full tray.

Pupal survival and sex ratio were evaluated by incubating 100 pupae in a lidded cup until all adults had emerged, roughly 72 hr after first adult emergence. Dead pupae and adult sex ratios were counted from each cage.

Biological yields were evaluated by the total egg mass produced per cage, total pupae mass of pupae produced, the average individual pupa mass per cage, estimated number of pupae produced per cage, and average adult male mass at 4 days old. Average pupae volume was estimated by the weight of 20 pupae and adult male mass by the weight of 5 males.

### Mating and longevity evaluation

Mating capability was evaluated using both non-competitive and competitive mating tests. Non-competitive tests were completed by placing 5 males with 15 virgin females in a cage and allowing free mating for 4 hours. All flies were 5 days old. After 4 hr, males were removed. Female mating status was determined by extraction of their spermatheca and visually confirming the presence of sperm. For DR6, DR7, and J-06 males, female J-06 were used.

Competitive mating was evaluated by placing 10 DR6 or DR7 x EF1 double homozygous strain males with 10 J-06 males and 10 J-06 females for 20 hr. After mating, males were removed and females induced to oviposit individually. Egg masses produced were allowed to hatch and the larvae were observed for fluorescence and sterility. Sterile eggs were counted as non-mated.

Longevity was evaluated per strain by placing 25 pupae in two cages and provided food. Daily mortality of males was counted per cage. Food was replaced every 7 days and water was not replaced through the duration of the test.

### Statistics

Statistical analyses were completed using the R statistics program (R Core Team 2016). Biological yield comparisons between +Dox and -Dox treatments and between the full and reduced Dox feeding regimens was completed using analysis of varaiance (ANOVA) with TukeyHSD posthoc test for all parameters with the exception of male mass where a Student’s T-test was used. Pairwise comparisons of survival percent were done using the Wilcoxon signed-rank test. Differences in non-competitive and competitive mating success were evaluated using a pairwise Fisher’s Exact Test with R package rstatix (ver. 0.7.2; Kassambara, 2023). For competitive mating assays comparisons used counts from successful matings only, females that did not lay eggs or laid infertile eggs were excluded from totals. Longevity analysis was completed using the survival package (ver. 3.7; Therneau, 2023) by Cox proportional hazards regression model with multiple comparisons of means by Tukey contrasts and plotted with ggplot2 (ver. 3.5.1; Wickham, 2016).

## Results

### Strain creation

Transgenic screwworm strains developed in this study were obtained through microinjection of plasmid DNA into embryos from the production strain Jamaica-06 (J-06), the strain produced by the screwworm sterile fly production plant, COPEG, from 2006 to 2024. EF1 effector lines were developed previously also with the J-06 strain (Concha et al. 2020). Microinjection of DR6 plasmids resulted in 6 stable lines. Out of these lines, DR6#2 and DR6#5 were selected for further characterization as they produced the most stable lines with consistent female mortality when crossed to effector lines EF1#16. Initial injections of DR7 plasmids resulted in one stable strain, DR7#19, and subsequent injections of DR7 produced 37 independent transgenic lines. These secondary DR7 lines were reduced to 7 lines either by line failure or random selection to reduce workload. DR7#19 was crossed with EF1#19 to produce a stable line, while DR7#A6 was crossed to EF1#16. Both EF1lines produced similar levels of female lethality in previous strain crossings.

Two component DR6 or DR7 with EF1 lines were developed using Tc to suppress female mortality, but neither functioned as predicted. DR6 x EF1 strains were stable, but in the absence of Tc female death was delayed until pupal development, rather than the predicted embryo lethality. DR7 x EF1 lines were highly unstable and frequently resulted in infertile females, low larval survival, or high rates of pupal mortality even when reared with Tc. Replacing Tc with Dox improved DR7 x EF1 line stability significantly and resulted in lines suitable for further analysis and assessment of suitability for mass-rearing (Arp et al. 2024).

### Gene expression analysis

Verification of driver function was completed by qPCR and relative quantification of the tTAo mRNA (Figure 2). In all lines, expression of *tTAo* RNA was greatest in 2-3 hr embryos. Unexpectedly, tTA was also expressed at high levels in pupae in three of the four strains. In the DR6#A6 x EF1#16 strain, tTA expression in pupae was low. In all strains there was little tTA detected in the 2^nd^ instar larvae or adult samples. Expression of tTA was not equal between drivers DR6 and DR7. In DR6 strains, tTA was expressed more equally in all three embryo stages tested and expression in 2-3 hr embryos was approximately double the other larval stages. In DR7 strains, tTA was seven times more expressed in the 2-3 hr embryos than DR6 strains. The high level of tTA RNA in very early 0-1 hr embryos in the DR6 strains suggests significant maternal expression and RNA deposition into the developing eggs.

**Figure 2.**
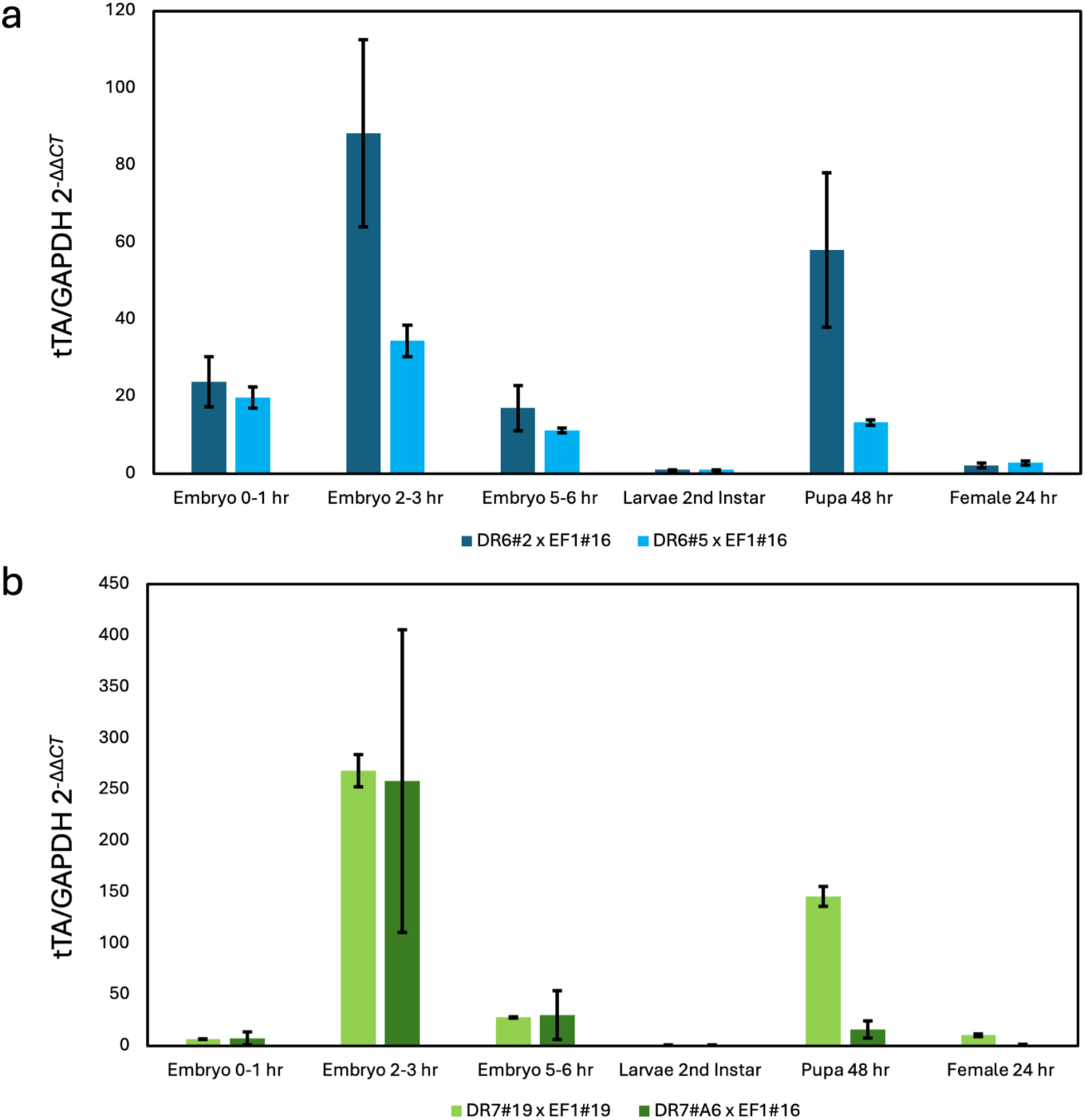
Relative expression of *tTAo* RNA during development. a) DR6 strains and b) DR7 strains. RNA was isolated from precellular (0-1 hr), cellular (2-3 hr) and late stage embryos (5-6 hr) as well as second instar, pupae (48 hr) and young adult females (24 hr). RNA levels are in relation to GAPDH with larval 2^nd^ instar samples used as the calibrator. Mean from three biological replicates and three technical replicates per reaction with standard error shown.

### Female stage lethality

Female survival observations were made by comparing survival rates between +Dox and -Dox conditions under both Dox feeding regimes (Figure 3, Table S1). The expected outcome under the Dox-feeding condition was that half of the embryos were female and would die before emerging. Consequently, larval counts would reflect embryo hatch rates though development.

**Figure 3.**
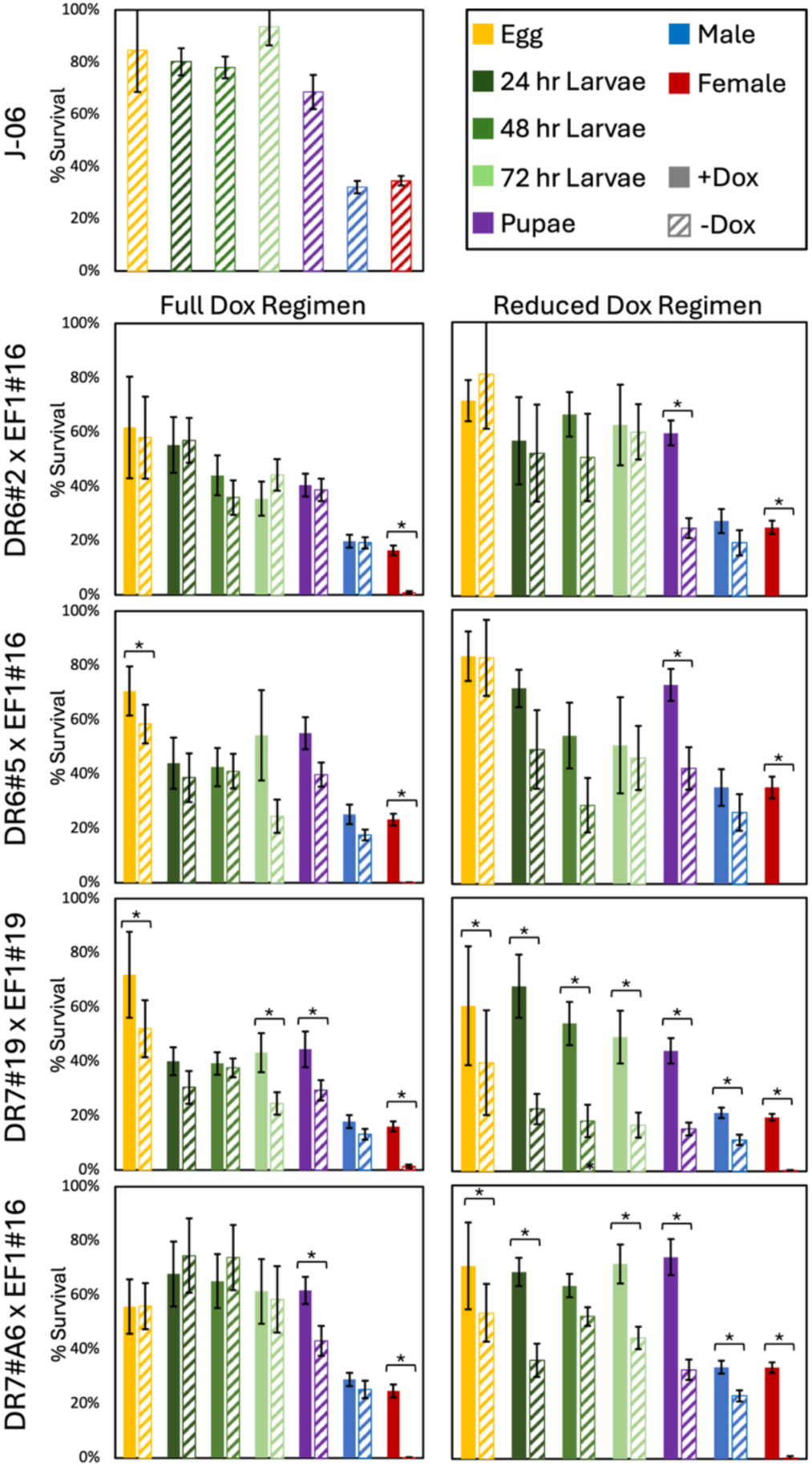
Staged lethality of the wild type (J-06) and transgenic sexing strains. Graphs show the estimated mean survival rate through development for each line on diet with (+Dox) or without (-Dox) Dox. Survival estimates included are egg hatch, 1^st^, 2^nd^, and 3^rd^ instar larvae, pupae, and adult male and female. Both full and reduced Dox feeding regimens are shown. Standard error of the mean is displayed on bars. Significant (p < 0.05) pairwise comparisons between +Dox and -Dox treatments are indicated with “*” and brackets (Table S1).

Both DR6 strains showed few significant differences in embryo or larval survival between the +Dox and -Dox treatments under the full Dox feeding regimen, implying that females were not dying before pupation (Figure 3, Table 1). On the -Dox treatment roughly half of the pupae were male while the remaining pupae did not survive, suggesting that the remaining unhatched pupae were female. On the reduced Dox regimen both DR6 strains had no significant differences in egg hatch rates and larval counts between the +Dox and -Dox treatments, but a roughly 50% reduction in pupae produced on the -Dox treatment, suggesting that females on the reduced Dox regimen died in late larval development prior to pupation.

**Table 1.**
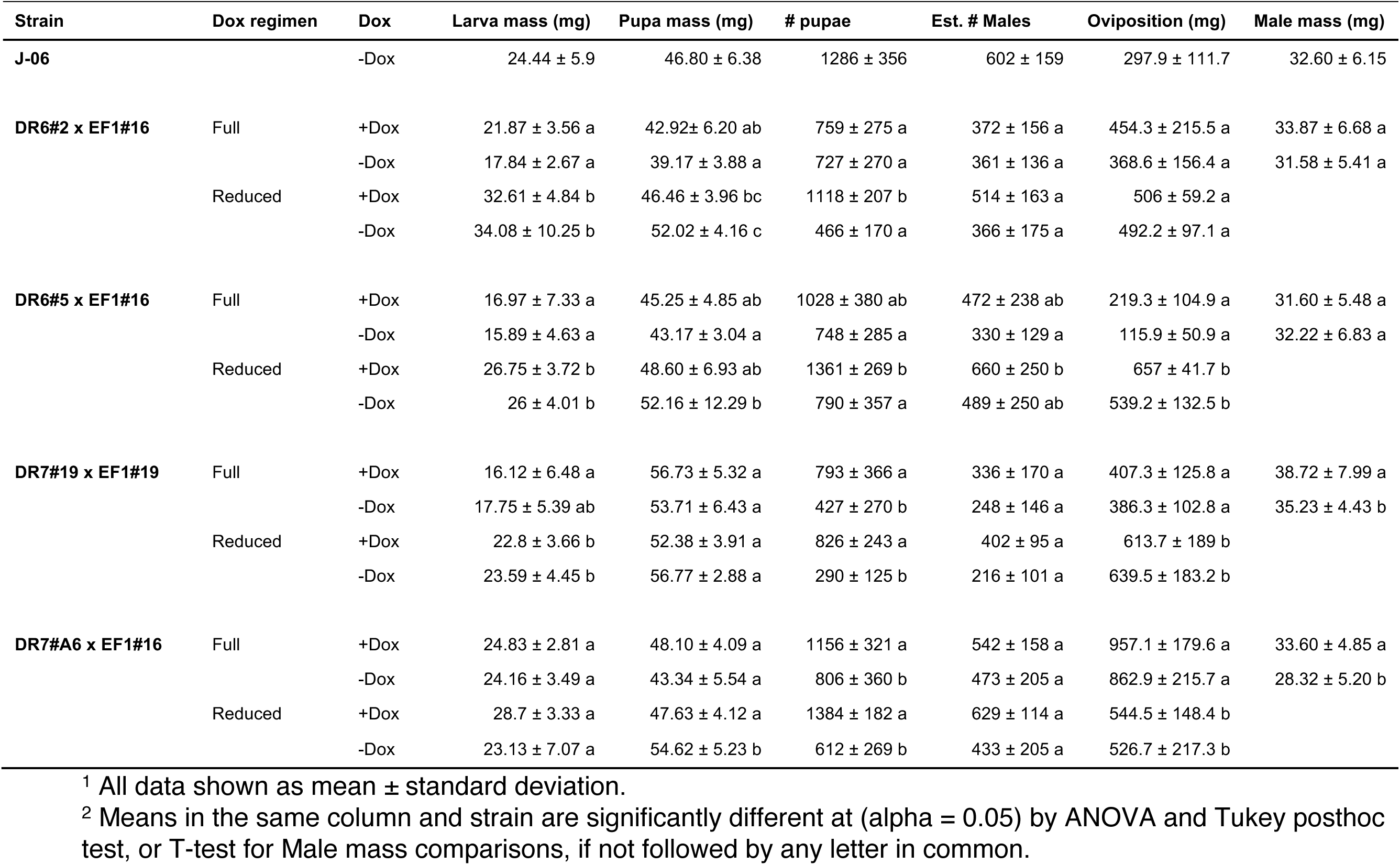
Biological performance metrics of all strains.

The DR7#19 x EF1#19 strain had reduced late embryo survival in the -Dox condition under the full Dox regimen, the reduced Dox regimen maintained early reductions in survival through larval development, implying the reduction in survival occurred in the embryo or first 24 hr of larval development and not during larval development. DR7#A6 x EF1#16 did not have apparent reductions in survival in the - Dox treatment under the full Dox regimen but did have significanly reduced survival starting from embryos, with the exception of 48 hr observations, under the reduced Dox regimen. Both DR7 strains showed a decrease in male survival on the reduced Dox regimen compared to the +Dox controls. On the Full Dox regimen a few females were obtained from the DR7#19 x EF1#19 strain but none survived under the reduced Dox regimen. Overall, these survival observations indicate that both DR6 strains are late larval or pupae lethal and DR7 strains are embryo or early larval lethal if given reduced dosages of Dox. In all treatments, females emerging from pupae in the -Dox treatment were malformed and not reproductively viable.

### Comparison of biological yields of the transgenic sexing strains relative to J-06

Overall, both DR6 and DR7 with EF1 strains were similar in larvae, pupae, and adult size with the production strain J-06 (Table 1). All statistical comparisons of biological size are included in Table S2. Inclusion of Dox in the diet and Dox regimen influenced strain performance, with regimen having the most significant impacts on biological yield metrics (Table 1). In all TSSs but DR7#A6 x EF1#16, 72hr larvae were larger on the reduced Dox regimen and were not impacted by the presence or absence of Dox. Likewise, the reduced Dox regimen also produced larger pupae in all strains but DR7#19 x EF1#19.

For all lines, the +Dox treatment is expected to produce double the number of pupae as the -Dox treatment. In the full Dox regimen, more pupae were produced in the +Dox treatment for all lines except DR6#2 x EF1#16 which produced an equal number of pupae in the + and - Dox treatments. The reduced Dox regimen produced a more expected reduction by approximately half for all strains, including DR6#2 x EF1#16. Although the number of pupae is expected to reduce, the number of surviving adult males should remain the same between the Dox treatment and feeding regimens. On the full dox regimen, there were no significant differences between the number males obtained between the + and - Dox treatments for three of the strains. The exception was DR6#2 x EF1#16 with a mean number of males that was lower in the -Dox treatment. In the reduced Dox treatment, there was a significant reduction in the number of males produced by DR7#19 x EF1#19 (46% decrease) and DR7#A6 x EF1#16 (31% decrease) in comparison with the full Dox regimen.

Male mass was only observed in the full Dox regimen. No differeneces were observed between treatments for both DR6 strains, but both DR7 strains were smaller on the -Dox treatment. Oviposition mass produced by the females on the reduced Dox regimen increased for both DR6#5 x EF1#16 and DR7#19 x EF1#19 and was reduced in DR7#A6 x EF1#16.

### Mating and longevity

Mating capability and longevity are important parameters for predicting performance of males in the field. In the non-competitive mating evaluation (Figure 4a), both DR6#2 x EF1#16 (p < 0.001) and DR6#5 x EF1#16 (p < 0.001) mated with significantly less females during the access period than J-06. However, DR7#19 x EF1#19 (*p* = 0.81) and DR7#A6 x EF1#16 (*p* = 0.69) were comparable with J-06, mating with at least an equal number of females in the limited time available for mating. When males of J-06 and DR6 or DR7 TSSs were put into competition for a limited number of J-06 females (Figure 4b), J-06 females mated significantly more with J-06 males than DR6#2 x EF1#16 (Χ^2^ = 44.31, df = 1, *p* < 0.001), DR6#5 x EF1#16 (Χ^2^ = 27.56, df = 1, p < 0.001), DR7#19 x EF1#19 (Χ^2^ = 8.257, df = 1, p < 0.004), or DR7#A6 x EF1#16 (Χ^2^ = 26.79, df = 1, p < 0.001) males. Longevity of TSS males was observed in the -Dox treatment of the full Dox feeding regimen (Figure 4c). The only TTS with equal longevity to J-06 was DR6#2 x EF1#16 (z-value = -0.323, *p* = 0.998). All other TSS had significantly reduced longevity than J-06 (p < 0.001). DR6#5 x EF1#16 had higher mortality in the first 10 days than other strains but was equal to J-06 in 50% survival time.

**Figure 4.**
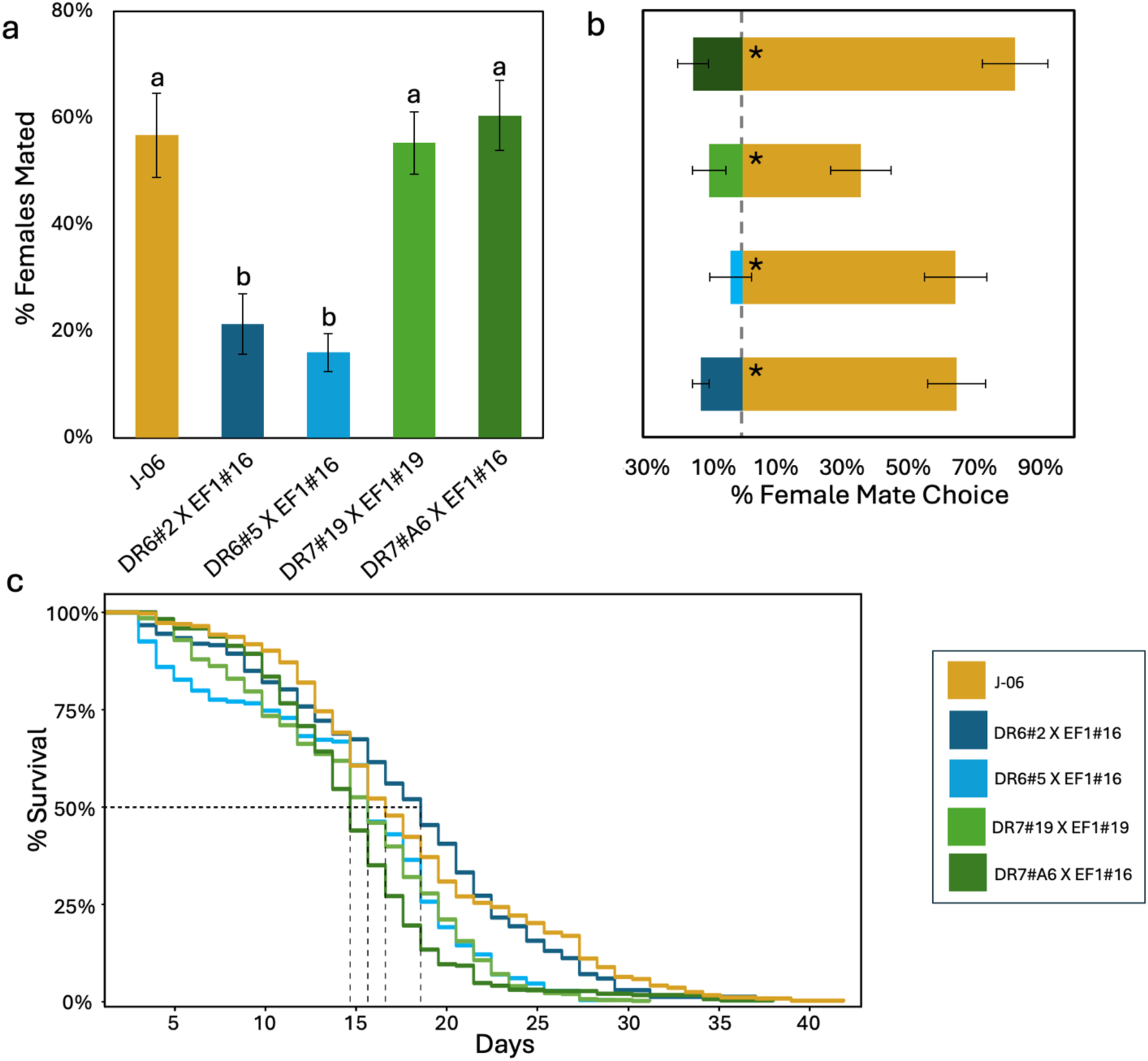
Mating success, mating competitiveness and longevity of TSS males. a) male mating success in non-competitive arenas. Statistical differences are indicated with letters. b) female mate choice under competitive ratios of 10 TSS males : 10 wildtype (J-06) males : 10 wild type females. Significant differences in mate selection are indicated by “*”. c) survival probability of adult males from Dox-rearing. For panels a and b data shown as mean with standard error.

## Discussion

Insect control programs that utilize SIT can be much more effective if the sterile insects are only males (Rendon et al. 2004). SIT programs utilize different methods to separate sexes such as dimorphic features like pupa size, while others have developed strains with heritable sex-linked traits. Screwworm do not have any sex distinguishable features before the adult stage and no classical sex-separating strains have ever been developed, so transgenic technologies were deployed to create conditional female lethal strains. Previously developed screwworm TSSs were able to achieve high levels of reliable female lethality, but lethality occurred too late in development to provide cost reductions (Concha et al. 2016) or required antibiotic regimens with potential to produce infertile colony females or rescue female embryos in desired all-male generations if applied incorrectly in mass-rearing (Concha et al. 2020). Updated driver strains, DR6 and DR7, with predicted expression limited to early embryos were developed in *L. cuprina* and found to be effective at eliminating the need for adult provisioning with Tc and likely suitable for use in a mass-rearing system (Yan et al. 2020).

When evaluated in screwworm, drivers DR6 and DR7 functioned as expected with expression peaking in 2-3 hr embryos but with key differences to *L. cuprina* TSSs carrying these drivers. *L. cuprina* DR6 and DR7 lines expressed tTA RNA at similar levels but screwworm DR7 strains were over 4 times more active than DR6 in 2-3 hr embryos. This is likely because the *CmCG14427* promoter used for DR7 was from *C. macellaria*, which is much more closely related to *C. hominivorax* than *L. cuprina* (Yan et al. 2020). This suggests that the gene promoters from *C. hominivorax* cellularization genes identified previously (Scott et al. 2020) could also drive high levels of tTA RNA expression. Not seen in *L. cuprina*, both drivers had elevated tTA expression in the pupae of screwworm DR6 lines and one of the DR7 lines. Differences were also observed between the two evaluated DR7 lines, with DR7#A6 having high variability in tTA RNA production between replicates from embryos. Yan et al. (2020) observed that the endogenous *Lcnullo* and *LcCG14427* genes had a much greater difference in expression level between early embryo and adult stages (about 1000x) compared to the DR6 and DR7 lines (about 10 times). They suggested this could be the result of epigenetic control mechanisms that suppress the *Lcnullo* and *LcCG14427* genes in adults in their normal chromosomal location but not in transgenes driven by the gene promoters. These missing regulatory elements could explain the pupal expression in the screwworm DR6 and DR7 lines and the elevated expression observed in DR7 lines.

Differences in screwworm DR6 and DR7 expression was reflected in strain health and stage lethality. On the reduced Dox feeding regimen, both DR6 TSSs showed 100% female lethality in late larval development, while for the DR7 TSSs females appeared to die at the embryo and early larval stages. Although *tTAo* expression in the DR6 lines was highest in 2-3 hr old embryos, the delayed female mortality observed in the TSSs indicates insufficient production of Lshid^Ala2^ to result in immediate mortality but causing issues in later development. Although the DR7 TSSs were more effective at eliminating females early in development this came at a cost of a reduction in survival and total production. The DR7 TSSs produced significantly fewer males on the reduced Dox regimen relative to +Dox (46%, 31% decrease) and the J-06 wild type (64%, 28% decrease). As overexpression of tTA can be lethal or reduce survival (Concha et al. 2016), it is possible that the high levels of expression of *tTAo* in the DR7 lines is negatively impacting strain performance. Further, low levels of the female mode of splicing of the *Chtra* intron in *Lshid^Ala2^* would lead to “leaky” expression in some males resulting in reduced strain performance. Yan et al. (2020) offered both explanations for why two of the four *L. cuprina* DR6/DR7 TSSs showed low pupal eclosion frequencies on diet without tetracycline. As the TSSs described here did have differences in *tTAo* expression, it is difficult to discern if reduced strain performance is due to leaky expression of *Lshid^Ala2^*, tTA toxicity, or a combination. Overall, the most promising TSS produced in this study was DR7.

Proper dosage of Dox was an important factor in strain function. Previous work with the TD1 all-in-one TSS using the *bottleneck* promoter had complete female lethality when larva of the parent colony was fed diet with 25 ug/ml Dox or 200 ug/ml Tc (Concha et al. 2020; Arp et al. 2024). Rearing DR7 lines with 200 ug/ml Tc resulted in inconsistent performance often causing female infertility or high mortality throughout development (unpublished data). Changing the sexing system suppressor from Tc to Dox aided in stabilizing strain performance but prevented female mortality until the late larvae or pupal stages. The greater bioavailability of Dox likely explains why it was more effective in switching off the female lethal system and facilitating strain maintenance. Removing Dox from the last larval feeding shifted female mortality to embryo or early larval development. The short active window early in embryonic development of both promoters was likely interrupted by maternal transfer of Dox due to receiving constant diet additions containing Dox prior to oviposition.

The characterization of screwworm TSSs driven by promoters of *Lcnullo* and *CmCg14427* found significant differences in gene expression resulted in complex interactions. High transgene expression observed in DR7 lines resulted in embryo female mortality, but reduced overall strain performance and production capacity, while lower gene expression observed in DR6 lines resulted in higher productivity but late female mortality. The DR7 lines also required the use of Dox to improve strain reliability and stability, but the stability of Dox resulted in female rescue through maternal transfer. The DR6#A6 x EF1#16 strain was the most promising TSS made in this study. The strain shows 100% female lethality in early development on the reduced Dox regimen, males were comparable to J-06 in the mating success assays and male production was only about 25% lower than J-06. However, male longevity was significantly reduced and males were outcompeted by J-06. Consequently, TSSs made earlier such as TD1 (Concha et al. 2020) would be a higher priority for future evaluation under mass rearing conditions and in open field tests. Nevertheless, the DR6 and DR7 TSSs provided a challenging testing ground to evaluate how driver constructs function in different genetic backgrounds and how optimizing antibiotic dosages for transgenic strains is essential to maintaining them, especially when being considered for use in a factory setting where greater tolerances are necessary to ensure constant production.

## Acknowledgements

The authors would like to thank Rosaura Sánchez, Nicolas Mendoza, Domitildo Martínez, Hermogenez González, Odilis Rodríguez, Kenneth Castillo, and Joel Sanchez for their contributions to rearing strains. This research used resources provided by ARS project number 3094-32000-041-00D and through agreements between NC State University with COPEG (01-15) and USDA-APHIS (AP19IS000000C001). Mention of trade names or commercial products in this publication is solely for the purpose of providing specific information and does not imply recommendation or endorsement by the USDA. This research was supported in part by the U.S. Department of Agriculture, Agricultural Research Service. USDA is an equal opportunity provider and employer.

## Author Contributions

A.P.A and M.J.S. designed the study; A.P.A, and C. C. conducted embryo microinjections and strain crossing. Y.Y and M.J.S designed gene constructs. A.V.L., M.V., G.Q., and A.S. maintained fly strains and conducted biological assays; A.P.A., A.V.L., and A.M isolation of RNA and expression analysis. A.P.A analyzed raw data and conducted statistical analyses. M.J.S. obtained funding and provided supervision, A.P.A. and M.J.S. wrote the manuscript with input from A.V.L., A.S., G.Q., M.V., C.C., and Y.Y.

## Supplemental Tables

**Table S1.** Pairwise satistsical comparisons of stage specific survival.

**Table S2.** Pairwise satistsical comparisons of biological performance metrics.

